# FOXP3+ Regulatory T Cells Require TBET to Regulate Activated CD8+ T Cells During Recovery from Influenza Infection

**DOI:** 10.1101/2024.05.30.596295

**Authors:** Nurbek Mambetsariev, Manuel A. Torres Acosta, Qianli Liu, Carla P. Reyes Flores, Anthony M. Joudi, Kathryn A. Helmin, Jonathan K. Gurkan, Elizabeth M. Steinert, Luisa Morales-Nebreda, Benjamin D. Singer

**Author notes:** To whom correspondence should be addressed: Benjamin D. Singer, MD, Division of Pulmonary and Critical Care Medicine, Department of Medicine Department of Biochemistry and Molecular Genetics, Northwestern University Feinberg School of Medicine, 303 E. Superior St., Simpson Querrey 5^th^ Floor Chicago, IL 60611 USA. These authors contributed equally (Nurbek Mambetsariev, Manuel A. Torres Acosta). Competing Interest Statement: NM is currently an employee and owns stock in Vertex Pharmaceuticals. BDS holds United States Patent No. US 10,905,706 B2, Compositions and Methods to Accelerate Resolution of Acute Lung Inflammation, and serves on the Scientific Advisory Board of Zoe Biosciences. The other authors have no competing interests to declare.

## Abstract

FOXP3+ regulatory T (Treg) cells are necessary to coordinate resolution of lung inflammation and a return to homeostasis after respiratory viral infections, but the specific molecular requirements for these functions and the cell types governed by Treg cells remain unclear. This question holds significance as clinical trials of Treg cell transfer therapy for respiratory viral infection are being planned and executed. Here, we report causal experiments in mice determining that Treg cells are necessary to control the numbers of activated CD8+ T cells during recovery from influenza infection. Using a genetic strategy paired with adoptive transfer techniques, we determined that Treg cells require the transcription factor TBET to regulate these potentially pro-inflammatory CD8+ T cells. Surprisingly, we found that Treg cells are dispensable for the generation of CD8+ lung tissue resident-memory T (Trm) cells yet similarly influence the transcriptional programming of CD8+ Trm and activated T cells. Our study highlights the role of Treg cells in regulating the CD8+ T cell response during recovery from influenza infection.

---

Tissue resident memory CD8+ T (Trm) cells are generated at the site of an acute infection from a pool of antigen-specific infiltrating lymphocytes (1). In some tissues, Trm cells persist over time, although the influenza-specific Trm compartment dramatically contracts in the lungs of young mice over a period of months following influenza infection (2). Most Trm cells sequentially express the activation marker CD69 followed by integrin alpha E (CD103), and this combination has been used to detect Trm cells (1). Following influenza infection, CD4+ T cells are required for the formation of lung-resident Trm cells (3). Emerging evidence has implicated CD4+ FOXP3+ regulatory T (Treg) cells in Trm development in the lungs and other tissues (4, 5). Following viral clearance, Treg cells resolve inflammation to promote recovery from influenza (6), but their effect on the Trm compartment during recovery from influenza is unknown. We hypothesized that Treg cells regulate the dynamics of Trm cells in the lung following influenza infection.

To determine whether Treg cells regulate Trm abundance, we intra-tracheally inoculated *Foxp3*^*DTR*^ mice with a sublethal dose of influenza virus. Following viral clearance (7), mice were treated with diphtheria toxin (DT) to deplete Treg cells during days 14 to 22 post-inoculation (**Figure 1A**) – a time window following peak recruitment of CD8+ T cells to the lungs (8). As expected, Treg cells were reduced in the lungs of DT-treated mice at days 22 and day 28 post-inoculation but recovered by day 45 (**Figure 2A**). The frequency of lung CD8+ CD69+ CD103+ Trm cells was reduced in the absence of Treg cells at day 28 (**Figure 2B**), but their absolute count in the lungs was not affected through day 45 (**Figure 1B**). In contrast, CD8+ T cells that expressed only CD69 (i.e., activated CD8+ T cells) were higher in both frequency (**Figure 2C**) and number through day 45 in the Treg cell-deficient group (**Figure 1C**). CD8+ T cells lacking CD69 and CD103 expression (double-negative cells) were also greater in number in the absence of Treg cells (**Figure 2D**), but intravascular labeling revealed the pulmonary circulation as the location of the majority of these cells (**Figure 2E**). Collectively, these results indicate that, following influenza infection, Treg cells regulate the number of CD8+ CD69+ (activated) T cells, which accumulate to a greater degree in the absence of Treg cells. In contrast, Treg cells do not affect the number of cells in the canonical CD8+ CD69+ CD103+ Trm compartment.

**Figure 1.**
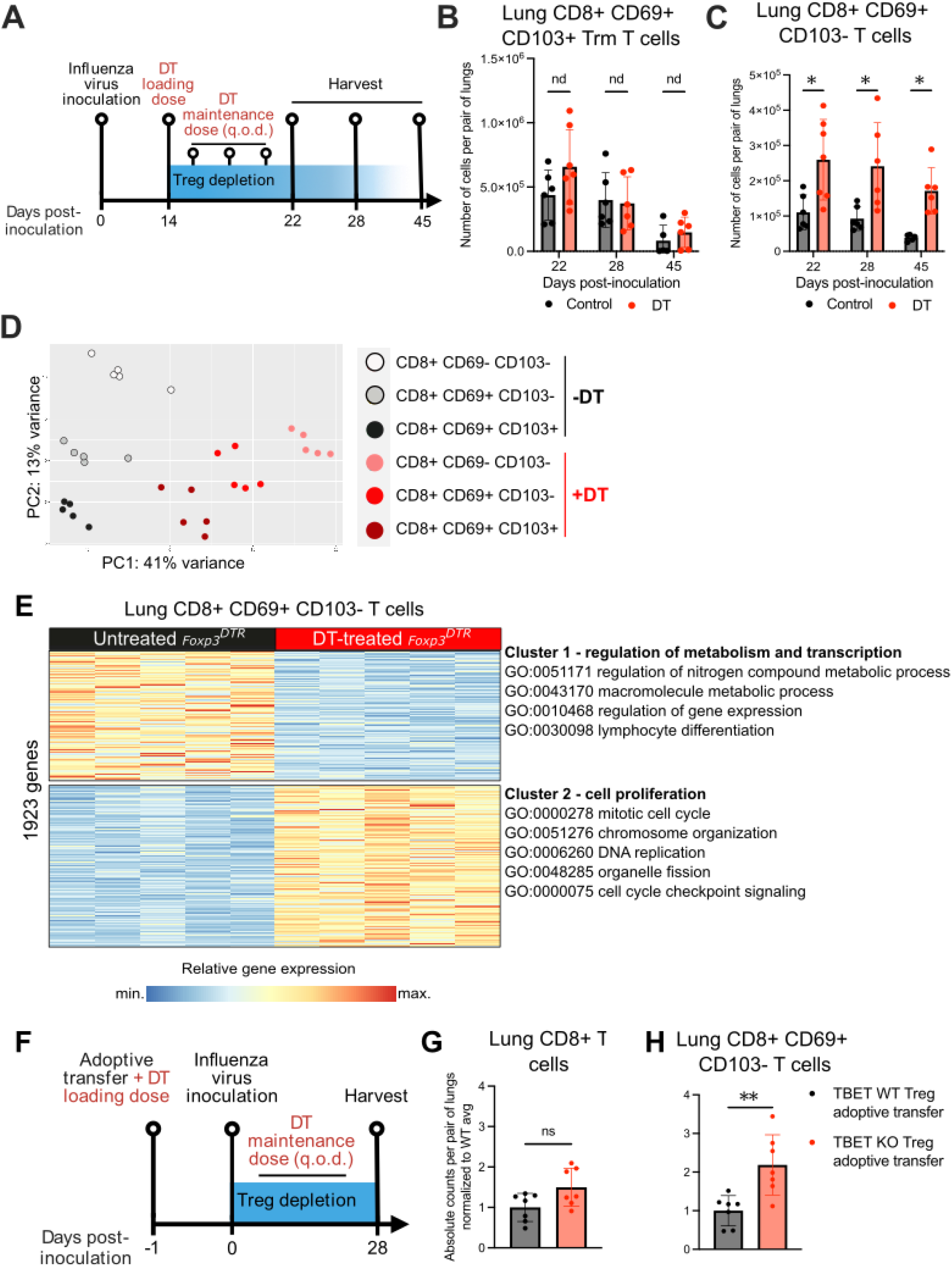
Treg cells regulate activated CD8+ cells in a TBET-dependent manner during recovery from influenza. (**A**) Experimental outline. *Foxp3*^*DTR*^ mice were intra-tracheally inoculated with 6 plaque-forming units of influenza A/WSN/33 H1N1 virus. At day 14 post-inoculation, a group of inoculated *Foxp3*^*DTR*^ mice received a diphtheria toxin (DT) loading dose (50 μg/kg) and three DT maintenance doses (10 μg/kg) every other day (q.o.d.). Lung CD8+ T cell subsets were quantified in DT-untreated (control) and - treated *Foxp3*^*DTR*^ mice at days 22, 28, and 45 post-inoculation. (**B**) CD8+ CD69+ CD103+ Trm cell and (**C**) CD8+ CD69+ CD103- (activated) T cell absolute counts per pair of lungs at days 22, 28, and 45 post-inoculation in control and DT-treated *Foxp3*^*DTR*^ mice. nd no discovery, * q < 0.05, according to Mann-Whitney U test with two-stage linear step-up procedure of Benjamini, Krieger, and Yekutieli with Q = 5%. Summary plots show all data points with mean and SD from two independent experiments per time point. (**D**) Principal component analysis of RNA sequencing data generated with sorted double-negative (CD69-CD103-), activated (CD69+ CD103-), and Trm (CD69+ CD103+) CD8+ T cells from DT-untreated and -treated *Foxp3*^*DTR*^ mice at day 28 post-influenza virus inoculation. (**E**) K-means clustering of differentially expressed genes (q < 0.05) and selection of top GO processes from the comparison of activated CD8+ T cells sorted from DT-untreated and -treated *Foxp3*^*DTR*^ mice at day 28 post-influenza virus inoculation. (**F**) Experimental outline. *Foxp3*^*DTR*^ mice received either TBET-sufficient (WT) or -deficient (KO) Treg cells via retro-orbital adoptive transfer alongside an intra-peritoneal diphtheria toxin (DT) loading dose (50 μg/kg). These mice received a maintenance dose of DT (10 μg/kg) every other day (q.o.d.) thereafter. One day after adoptive transfer, *Foxp3*^*DTR*^ mice were intra-tracheally inoculated with 6 plaque-forming units of influenza A/WSN/33 H1N1 virus. Lung T cells were quantified on day 28 post-inoculation. (**G**) Absolute counts of total CD8+ cells and (**H**) CD8+ CD69+ CD103-T cells per pair of lungs at day 28 post-influenza virus inoculation in DT-treated *Foxp3*^*DTR*^ mice that received either TBET WT or KO Treg cells. Data points were normalized to the TBET WT Treg adoptive transfer group’s average. ns not significant, ** p < 0.01 according to Mann-Whitney U test.

**Figure 2.**
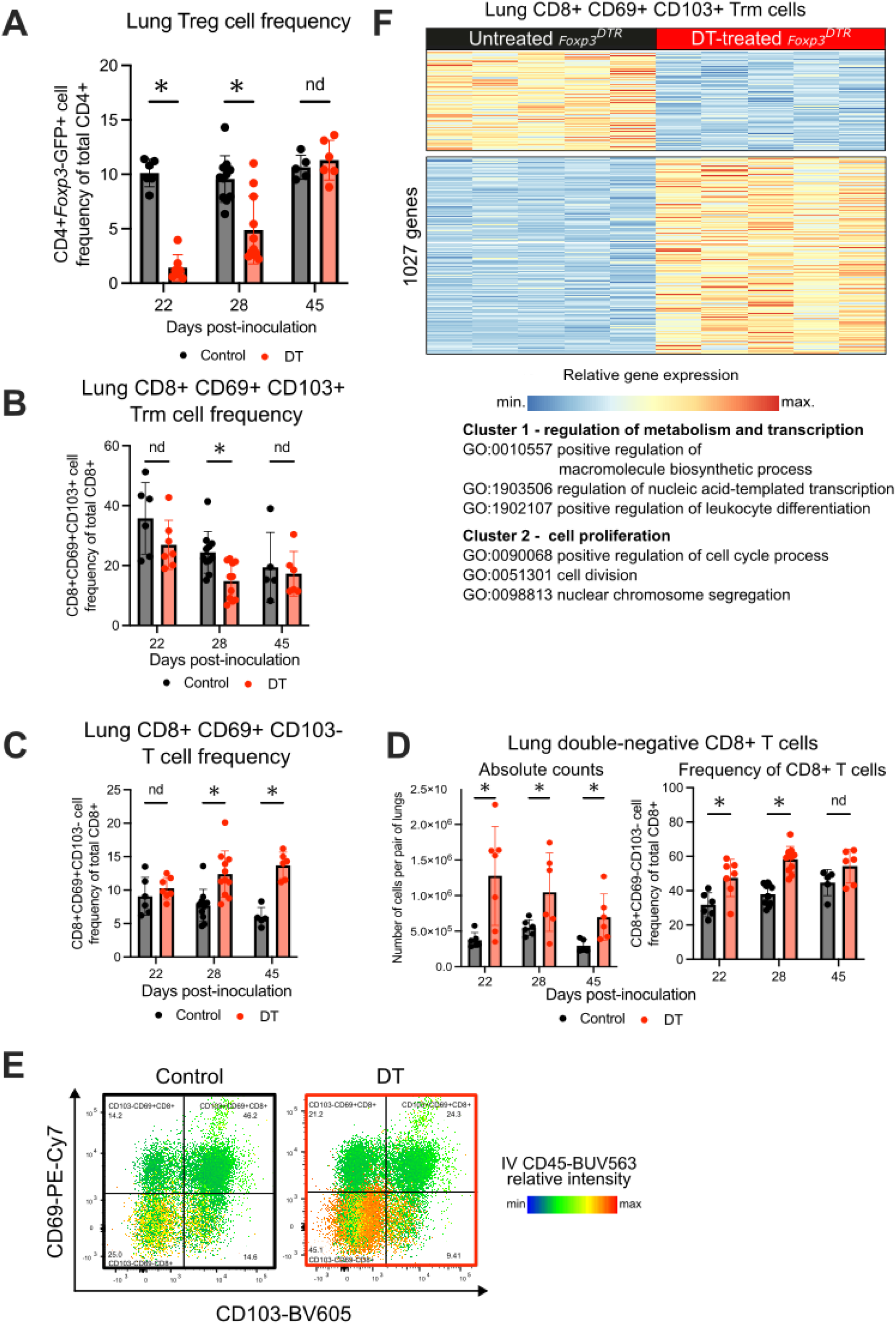
Characterization of T cell subsets during recovery from influenza in the absence of Treg cells. (**A**) Treg, (**B**) CD8+ CD69+ CD103+ Trm, and (**C**) CD8+ CD69+ CD103- (activated) T cell frequencies and (**D**) absolute count (left) and frequency (right) of double-negative CD8+ T cells in the lungs from the experiment outlined in Figure 1A. nd no discovery, * q < 0.05 according to Mann-Whitney U test with two-stage linear step-up procedure of Benjamini, Krieger, and Yekutieli with Q = 5%. Summary plots show all data points with mean and SD. (**E**) Representative heatmap statistic plot showing IV CD45-BUV563 signal intensity of lung CD8+ T cell subsets in control and DT-treated *Foxp3*^*DTR*^ mice that received 50 μg of CD45-BUV563 antibody retro-orbitally before harvest. (**F**) K-means clustering and selection of top GO processes from the comparison of CD8+ CD69+ CD103+ Trm T cells sorted from DT-untreated and -treated *Foxp3*^*DTR*^ mice at day 28 post-influenza virus inoculation.

We performed RNA sequencing of sorted CD8+ T cell subsets 28 days post-inoculation to ascertain whether Treg cells are a determinant of transcriptional programming in these subsets. Principal component analysis (PCA) of the RNA sequencing data segregated the DT-treated and -untreated groups along PC1, while the different CD8+ T cell subsets clustered along PC2 (**Figure 1D**). K-means clustering and GO process enrichment analysis of CD8+ CD69+ T (**Figure 1E**) and CD69+ CD103+ Trm (**Figure 2F**) cells revealed upregulation of genes involved in cellular proliferation and downregulation of metabolic and transcriptional genes in Treg cell-depleted mice, consistent with the persistence of the CD69+ (activated) subset when Treg cells were depleted (9).

FOXP3+ Treg cells can co-express T helper (Th) cell lineage-defining transcription factors to drive their recruitment, retention, and function in immunologically skewed microenvironments, in part by inducing expression of homing molecules (e.g., TBET-induced expression of CXCR3 in Th1-skewed microenvironments) (6). Because viral pneumonia elicits a Th1 response, we hypothesized that Treg cells require TBET (aka TBX21), the Th1-defining transcription factor, to regulate activated CD8+ T cells during recovery from viral pneumonia. Accordingly, we performed adoptive transfer of either TBET-sufficient or -deficient Treg cells into *Foxp3*^*DTR*^ mice treated with DT and quantified CD8+ T cell subsets in the lung on day 28 post-inoculation (**Figure 1F** and **Figure 3A**). Importantly, no significant difference in endogenous or total Treg cell levels was detected between mice receiving TBET WT or KO Treg cells (**Figure 3B-C**), and the majority of lung Treg cells in mice that received TBET KO Treg cells were CXCR3-negative, consistent with their TBET-negative status (**Figure 3D**). Adoptive transfer of TBET-deficient Treg cells yielded a significantly higher number of CD8+ CD69+ (activated) T cells but not CD8+ CD69+ CD103+ Trm cells relative to controls; total CD8+ T cells and double-negative CD8+ T cells were not significantly different between groups (**Figure 1G-H** and **Figure 3E-F**). These data suggest that Treg cells require TBET to control the numbers of activated CD8+ T cells during recovery from viral pneumonia.

**Figure 3.**
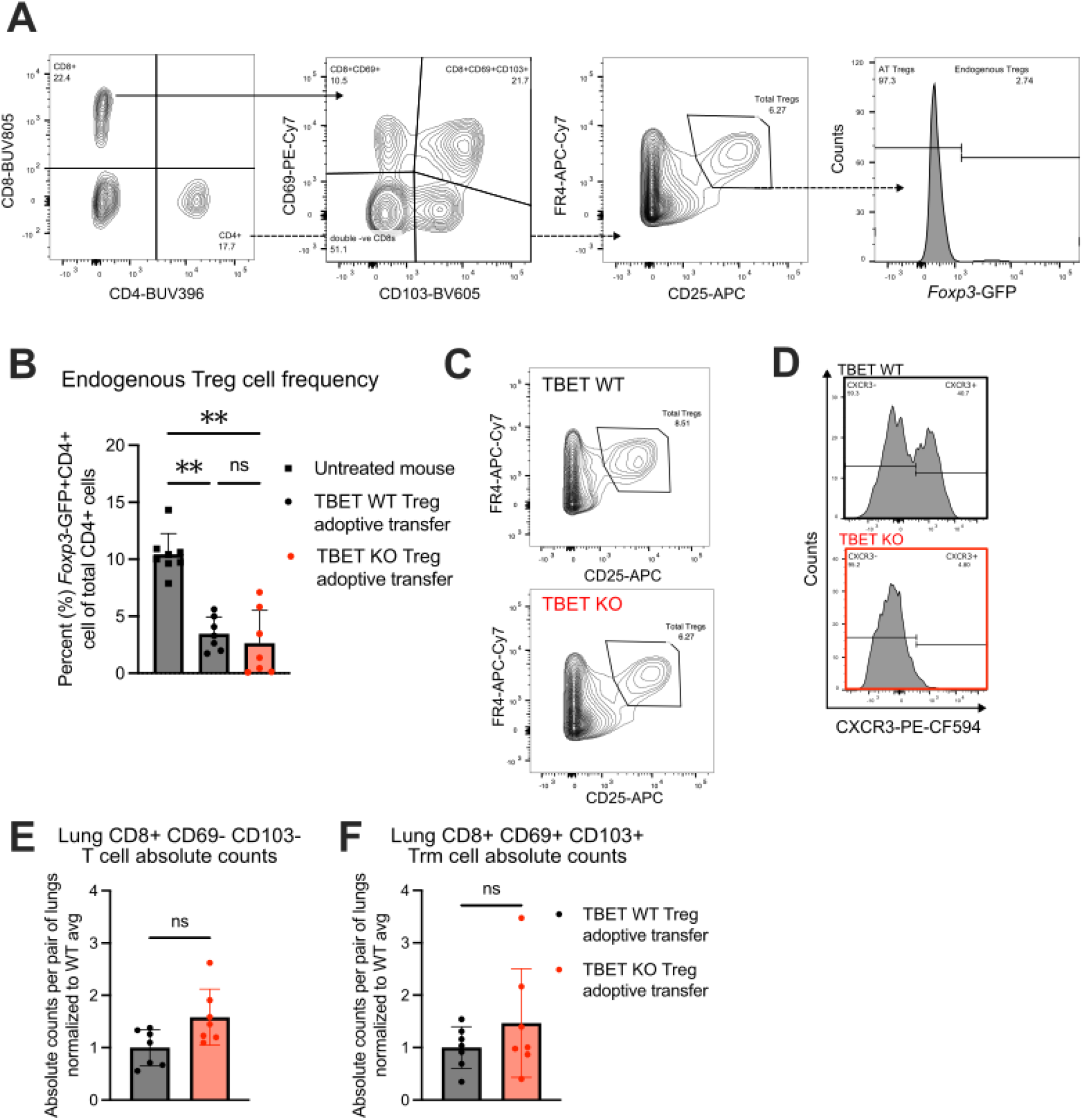
Necessity of Treg cell TBET expression on Treg, Trm, and other lung T cell numbers during influenza recovery. (**A**) Flow cytometry gating strategy for the quantification of T cell subsets, including CD8+ Trm cells (CD69+ CD103+), activated CD8+ T cells (CD69+ CD103-), double-negative CD8+ T cells (CD69-CD103-), endogenous Treg cells (CD4+ FR4+ CD25+ *Foxp3*-GFP+), and adoptive transfer Treg cells (CD4+ FR4+ CD25+ *Foxp3*-GFP-) in the experiment outlined in Figure 1F. (**B**) Frequency of *Foxp3*-GFP+ cells (endogenous Treg cells) of total CD4+ cells in DT-untreated *Foxp3*^*DTR*^ mice as well as DT-treated *Foxp3*^*DTR*^ mice that received either WT or TBET-deficient (KO) Treg cells. (**C**) Representative contour plots of total Treg frequency based on FR4 and CD25 staining in mice that received WT or TBET-deficient (KO) Treg cells. (**D**) Representative histograms of CXCR3+ Treg cell frequency of total Treg cells from DT-treated *Foxp3*^*DTR*^ mice that received either WT or TBET-deficient (KO) Treg cells. (**E**) Absolute counts of CD8+ CD69- CD103- (activated) T cells and (**F**) CD8+ CD69+ CD103+ Trm cells per pair of lungs at day 28 post-influenza virus inoculation from the experiment outlined in Figure 1F. Datapoints were normalized to the TBET WT Treg adoptive transfer group average. ns not significant, ** p < 0.01 according to Kruskal-Wallis test (B) or Mann-Whitney U test (E and F).

Our results demonstrate that TBET+ Treg cells regulate the numbers of activated CD8+ T cells expressing CD69 in the lung following influenza infection. Trm cells accumulated independently of Treg cells but mirrored transcriptional changes observed in activated CD8+ T cells following Treg cell depletion. Importantly, CD103 expression is not required for tissue residence of CD8+ T cells following influenza infection (10), and recent evidence suggests that expression of CD103 in Trm cells is dependent on the local microenvironment and TGFβ availability (11). Further studies are needed to test whether CD69 expression is sufficient to establish residency in activated CD8+ T cells persisting in the absence of Treg cells. Exploration of signals that promote CD103 expression in these cells can potentially improve protective memory in response to subsequent influenza infections or secondary pneumonias (12, 13). In contrast, long term persistence of Trm cells in the lungs promotes chronic pathology following influenza infection in old mice, and inhibiting their accumulation has therapeutic potential (14). Our study highlights the role of Treg cells in regulating the CD8+ T cell response during recovery from influenza infection.

## Materials and Methods

### Mice

All mouse procedures were approved by the Northwestern University IACUC under protocols IS00012519 and IS00017837. B6.129(Cg)-Foxp3tm3(Hbegf/GFP)Ayr/J mice (cat. no. 016958, referred to as *Foxp3*^*DTR*^ mice), B6.129S6-Tbx21tm1Glm/J mice (cat. no. 004648, referred to as TBET KO mice), and C57BL/6J mice (cat. no. 000664, referred to as Black 6 mice) were acquired from The Jackson Laboratory. All animals were genotyped using services provided by Transnetyx, Inc. with primers provided by The Jackson Laboratory.

### Flow cytometry and cell sorting

Single-cell suspensions of organ tissues, blood, tumors, or cultured cells were prepared and stained for flow cytometry analysis and sorting as described previously (7, 15). See **Table 1** for a list of antibodies used in this report. Cell counts of single-cell suspensions were obtained using a Cellometer with AO/PI staining (Nexcelom Bioscience cat. no. SD014-0106) before preparation for flow cytometry. Data acquisition for analysis was performed using a BD LSRFortessa or Symphony A5 instrument with FACSDiva software (BD). Cell sorting was performed using the 4-way purity setting on BD FACSAria SORP instruments with FACSDiva software or with a microfluidics MACSQuant Tyto sorter (Miltenyi). Analysis was performed with FlowJo version 10.9.0 software. Dead cells were excluded using a viability dye for analysis and sorting (16).

**Table 1.** Antibodies used for flow cytometry.

| Antigen/Reagent | Conjugate | Clone | Company | Catalog no. | Notes |
| --- | --- | --- | --- | --- | --- |
| CD45 | BUV563 | 30-F11 | BD Horizon | 612924 |  |
| CD4 | BUV395 | GK1.5 | BD Horizon | 563790 |  |
| CD8 | BV711 | 53-6.7 | BioLegend | 100747 |  |
| CD25 | APC | PC61.5 | Invitrogen | 2154040 |  |
| CD103 | BV605 | 2E7 | BioLegend | 121433 |  |
| CD69 | PE-Cy7 | H1.2F3 | BioLegend | 104512 |  |
| CXCR3 | PE/Dazzle594 | CXCR3-173 | BioLegend | 126534 |  |
| FR4 | APC-Cy7 | eBio12A5 | Invitrogen | 25-5445-82 |  |
| Fixable Viability Dye eFluor 506 | N/A | N/A | eBioscience | 65-0866-14 | Used according to manufacturer instructions |
| Absolute counting beads | N/A | N/A | Invitrogen | C36950 |  |

### Influenza A virus administration

Mice were anesthetized with isoflurane, intubated orally with an 18-gauge angiocatheter, and given two 25-μL aliquots of 3 plaque-forming units for a total of 6 plaque-forming units of influenza A/WSN/33 H1N1 virus in PBS through the catheter using a Hamilton syringe, 10 seconds apart (7). After each aliquot, the mice were placed on their right side and then left side for 10-15 seconds before recovery from anesthesia. At this dose of influenza, mortality does not occur and mice do not experience significant weight loss.

### Diphtheria toxin treatment and adoptive transfer of Treg cells to *Foxp3*^*DTR*^ mice

For the Treg cell depletion studies, *Foxp3*^*DTR*^ mice were intra-peritoneally injected with a diphtheria toxin (DT, List Biologicals cat. no. 150) loading dose (50 μg/kg) on day 14 post-influenza virus inoculation and given three maintenance doses (10 μg/kg) every other day thereafter to deplete endogenous Treg cells. For the adoptive transfer experiments, *Foxp3*^*DTR*^ mice received a DT loading dose one day before influenza virus inoculation as well as 1 x 10^6^ Treg cells via retro-orbital adoptive transfer from either Black 6 or TBET KO mice enriched using the Miltenyi Biotec CD4+ CD25+ Regulatory T Cell Isolation Kit (cat. no. 130-091-041) according to the manufacturer’s instructions. *Foxp3*^*DTR*^ mice receiving adoptive transfers were then given a DT maintenance dose every other day until tissue harvest.

### Lung tissue harvesting and processing

Influenza virus-infected mice were euthanized and slowly infused with HBSS through the right ventricle of the heart, clearing the pulmonary circulation of blood. Some mice received 50 μg of CD45-BUV563 antibody retro-orbitally 45 minutes before harvest to label intravascular leukocytes. The lungs were harvested and grossly homogenized with scissors in HBSS containing 2 mg of collagenase D (Sigma Aldrich cat. no. 11088866001) and 0.25 mg of DNase I (Sigma Aldrich cat. no. 10104159001) per mL, incubated for 45 minutes at room temperature, and then further homogenized using the mouse lung protocol of the Miltenyi OctoMACS tissue dissociator (m_lung_02). These procedures have been previously reported (7, 17).

### RNA-sequencing and analysis

Flow cytometry-sorted cells were lysed immediately after sorting with QIAGEN RLT Plus containing 1% β-mercaptoethanol and subjected to RNA isolation using the QIAGEN AllPrep Micro Kit as previously described (17). RNA-seq library preparation was performed using the SMARTer Stranded Total RNA-Seq Kit, version 2 (Takara cat. no. 634411). After sequencing, raw binary base call (BCL) files were converted to FASTQ files using bcl2fastq (Illumina). All FASTQ files were processed using the nf-core/RNA-seq pipeline version 3.9 implemented in Nextflow (18, 19) with Northwestern University Quest HPC (Genomic Nodes) configuration (nextflow run nf-core/rnaseq -profile nu_genomics --genome GRCm38). In short, lane-level reads were trimmed using trimGalore! 0.6.5, aligned to the GRCm38 reference genome using STAR 2.7.11, and quantified using Salmon. All samples showed satisfactory alignment rate (>60%). After quantification, differential expression analysis was performed in R version 4.2.0 using DEseq2 v1.38.3 (20). Sample genotype was used as an explanatory factor. Within-group homogeneity was first confirmed by principal component analysis (PCA) and no outliers were found. K-means clustering of differentially expressed genes (FDR q < 0.05) was performed using a previously published custom R function (21). In brief, k was first determined using the elbow plot and the kmeans function in R stats 3.6.2 (Hartigan– Wong method with 25 random sets and a maximum of 1,000 iterations) was used for k-means clustering. Samples were clustered using Ward’s method and a heatmap was generated using pheatmap version 1.0.12. Subsequent GO term enrichment was performed using topGO version 2.50.0 with Fisher’s exact test. org.Mm.eg.db version 3.16.0 and GO.db version 3.16.0 was included in GO terms enrichment analysis as references.

### Statistics

p-values and FDR q-values resulting from two-tailed tests were calculated using statistical tests stated in the figure legends using GraphPad Prism v10. Differences between groups with *p* or *q* values < 0.05 were considered statistically significant. Central tendency and error are displayed as mean ± standard deviation (SD) except as noted.

## Data availability

Raw and processed sequencing data will be made publicly available via the GEO repository pending peer-reviewed publication.

## Acknowledgements

NM is supported by NIH award T32AI083216. MATA is supported by NIH awards T32GM144295, T32HL076139, and F31HL162490. QL is supported by the David W. Cugell Fellowship and the Genomics Network (GeNe) Pilot Project Funding. CPRF is supported by T32HL076139. AMJ is supported by NIH award F32HL162418. LM-N is supported by NIH awards K08HL159356, U19AI135964, and the Parker B. Francis Opportunity Award. BDS is supported by NIH awards R01HL149883, R01HL153122, P01HL154998, P01AG049665, U19AI135964, and U19AI181102. We wish to acknowledge the Northwestern University Flow Cytometry Core Facility supported by CA060553; the BD FACSAria SORP system was purchased with the support of S10OD011996. We also wish to acknowledge the Northwestern University Metabolomics and Integrative Genomics Core. This research was supported in part through the computational resources and staff contributions provided by the Genomics Compute Cluster, which is jointly supported by the Feinberg School of Medicine, the Center for Genetic Medicine, and Feinberg’s Department of Biochemistry and Molecular Genetics, the Office of the Provost, the Office for Research, and Northwestern Information Technology. The Genomics Compute Cluster is part of Quest, Northwestern University’s high performance computing facility, with the purpose to advance research in genomics.

